# A Generalized Life-Motion Mechanism Supports Invariant Directional Coding of Local Biological Kinematics in Humans

**DOI:** 10.64898/2026.03.12.705481

**Authors:** Zhihan Gao, Lianzi Xing, Rui Wang, Yi Jiang

## Abstract

Humans can detect biological motion (BM) from sparse local kinematic cues, yet whether the visual system encodes directional information in a category-invariant manner remains unresolved. Here we combined a visual adaptation paradigm with computational modelling to reveal direction-sensitive neural mechanisms specialized for local biological kinematics. Adapting to a side-view scrambled point-light walker produced a robust repulsive aftereffect in the perceived direction of an intact human walker near the frontal view. Notably, this aftereffect generalized across different terrestrial vertebrates (pigeon, cat, dog) and across various actions (running, crawling, cycling), demonstrating a high degree of kinematic invariance. Crucially, the effect disappeared when biological kinematics were disrupted (inversion or removal of gravitational acceleration cues) or when the test stimulus was replaced with non-biological object motion. Individuals’ direction-discrimination abilities were also highly correlated across diverse local BM patterns, indicating a shared underlying mechanism. Drift-diffusion modeling further revealed that adaptation primarily altered the efficiency of sensory evidence accumulation rather than decision-level processes, and individual drift-rate changes strongly predicted the magnitude of perceptual aftereffects. These findings provide compelling evidence for a generalized, direction-sensitive neural system tuned to local biological kinematics, extending the life-motion detector theory and revealing a fundamental principle of biological motion perception.

**Significance Statement:** Humans possess an exceptional ability to detect and interpret the movements of living beings—an ability fundamental to survival, social interaction, and adaptive behavior. But how does the visual system extract “life motion” so adeptly amidst the complexity of natural scenes? Using visual adaptation and computational modelling, we provide direct evidence for direction-sensitive neural mechanisms specialized for local biological kinematics. These mechanisms operate robustly across species and actions, and remain selective for natural biological dynamics. Our findings advance the life-motion detector theory by demonstrating that the human visual system encodes directional information in a highly invariant manner, revealing a fundamental computational strategy for detecting animate agents.

## Introduction

Recognizing and deciphering the movement of biological entities is crucial for human survival and development. The human visual system is particularly sensitive to biological motion (BM), even from simple animations represented by a set of dots depicting the movement of the agent’s major joints (Johansson, 1973). Such point-light animations carry a wealth of biologically and socially relevant information, including moving direction, gender, identity, and emotional state (Giese & Poggio, 2003). Among these, moving direction is a fundamental attribute of BM, as locomotion is the most common activity in daily life and its heading direction usually conveys one’s intention. Observers can readily discriminate the moving direction of point-light walkers against dynamic visual noise, even those presented peripherally and incidentally (Troje & Chang, 2013).

How do we spot life motion so adeptly amidst the chaos of the natural world? Obviously, one can extract the moving patterns relying on a holistic spatiotemporal organization of body motion, a process known as global BM perception (Beintema et al., 2006). In addition to these global visual cues, accumulating evidence points to a distinct mechanism based on the kinematics of individual joints, termed local BM processing. Strikingly, observers retain their ability to discern moving direction even when exposed to scrambled point-light BM displays lacking coherent global structure (Troje & Westhoff, 2006). Such directional information can be effectively extracted at a glance as short as 100 ms (Chang & Troje, 2009a), serving as a significant feature to guide subsequent behaviors (Wang et al., 2010). Moreover, sensitivity to local BM signals extends to different species (e.g., cats) and human actions (e.g., running) (Chang & Troje, 2008, 2009a). This sensory capacity appears to be innate, as it is found emerging very early in life (Bardi et al., 2014) and impervious to learning experiences after birth (Chang & Troje, 2009b). Notably, local BM perception is severely impaired once the animation is presented upside down (Troje & Westhoff, 2006). Further investigation reveals that gravity-compatible dynamics, such as vertical acceleration patterns, play a crucial role in local BM processing (Chang & Troje, 2009a). Given these insights, it has been proposed that there might exist an intrinsic brain mechanism which exploits the characteristics of local BM signals to rapidly detect the presence of living entities, functioning as a “life motion detector” (Troje & Chang, 2023). Yet, the brain encoding mechanisms underlying local biological kinematics remain poorly understood.

In the current study, we combined a visual adaptation paradigm with a direction discrimination task to address two key questions: (1) whether there exist neural representations specific to moving directions of local BM; (2) whether this direction encoding generalizes across species or actions. Visual adaptation refers to the phenomenon whereby prolonged viewing of a visual stimulus reduces neural responses selective for the adapted feature, leading to altered perception. This adaptation paradigm, termed the “psychologist’s microelectrode”, has been widely used to probe selective neuronal representations across stimulus dimensions, from simple features such as orientation to complex properties like facial identity (Webster, 2015). Although recent work has demonstrated that the walking directions of conspecifics is represented in human visual system (Chen et al., 2023; Jackson & Blake, 2010), there is still limited evidence regarding whether the brain encodes directions of local motion signals. In Experiment 1, we exposed participants to a side view of scrambled human walker sequences and measured the perceived direction of an intact human walker near the front view after adaptation (**Figure 1**). If there are neural modules sensitive to different directions of local BMs, adaptation to one direction may induce inhibitory processes within these modules, thus biasing perceived direction away from the adapting stimulus and resulting in a repulsive perceptual aftereffect. To further examine whether direction information is coded for specific local BM cues, we compared adaptation aftereffects elicited by inanimate motions (e.g., inverted scrambled BM, unnatural scrambled BM, and object motion). In Experiments 2 and 3, we investigated whether this neural representation of direction is robust to life motion regardless of changes in low-level attributes. Specifically, we evaluated cross-category adaptation aftereffects when the adapting and test stimuli differed in species (e.g., adapting to scrambled pigeons, cats, or dogs) or action (e.g., adapting to scrambled runners, crawlers, or cyclists).

**Figure 1.**
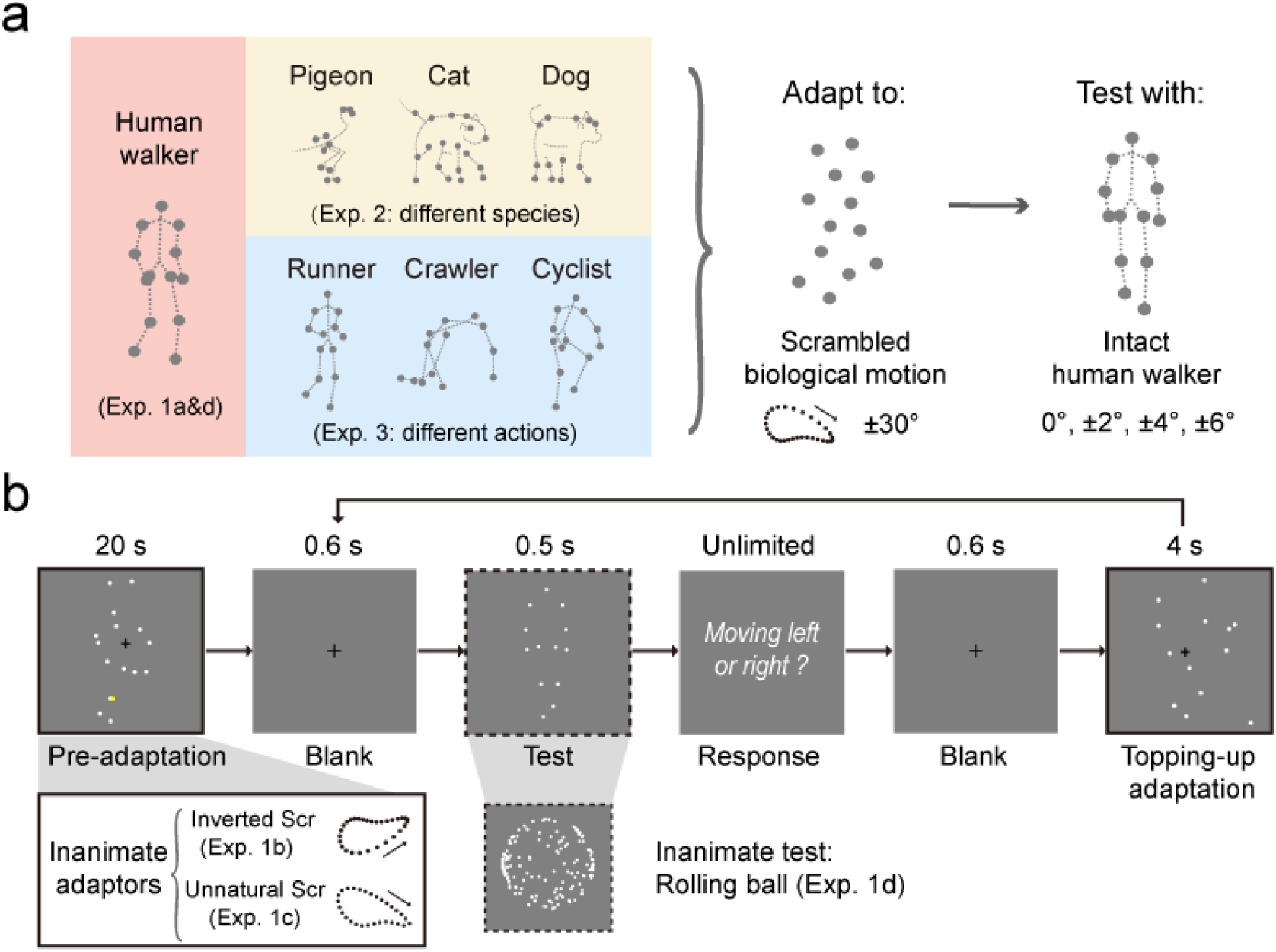
Stimuli and procedure. (**a**) Schematic of the point-light BM stimuli used across adaptation designs. Animate adaptors were spatially scrambled BM derived from human walkers (Experiment 1a), non-human animals (Experiment 2: pigeon, cat, dog), and other human locomotors (Experiment 3: runner, crawler, cyclist), shown from a side view. Intact human walker from a near front view served as the test stimulus. The dotted lines connecting critical joints of the agent were not present in the actual experiments and are shown here for illustration purposes only. (**b**) Experimental procedure. For the main adaptation paradigm (Experiments 1a, 2, and 3), each block began with a 20-s pre-adaptation to scrambled animate point-light sequences presented at ±30°side view. This was followed by a 0.6-s fixation and a 0.5-s test stimulus, an intact human walker from a near front view (0°, ±2°, ±4°, ±6°). Participants indicated the moving direction of the test stimulus (left or right). A 0.6-s intertrial interval and a 4-s topping-up adaptor preceded the next trial. During the pre- and topping-up adaptation, participants were asked to track the movement of the adapting stimulus and report any colour change. For control conditions, inverted (Experiment 1b) and unnatural (Experiment 1c) scrambled BM served as inanimate adaptors, featuring vertically inverted motion patterns and the removal of gravitational acceleration cues, respectively. In Experiment 1d, an upright scrambled BM served as the adaptor, while the test stimulus was replaced by a rolling ball.

## Results

### Experiment 1: Adaptation to scrambled human walkers

First, we tested whether adaptation to the spatially scrambled displays of a human walker from a specific viewpoint could affect the perceived direction of intact human walkers. As local BM processing is reported to be subject to an inversion effect (i.e., perception of local BM is severely impaired once the stimulus is presented upside down) (Troje & Westhoff, 2006), we contrasted the animate motions with inanimate motions that were generated by inverting the upright sequences. A group of participants underwent adaptation to upright scrambled point-light displays (PLDs) (Experiment 1a) as well as their inverted counterparts (Experiment 1b) in separate sessions. We observed a significant adaptation aftereffect when exposed to the upright scrambled PLDs of a human walker (**Figure 2a**, mean difference = 0.744°, *t* (15) = 3.153, *p* = 0.007, Cohen’s *d* = 0.788, CI = [0.241, 1.247], BF_10_ = 7.746). That is, after adaptation to a side view of local BM, participants tended to perceive an intact human walker near the frontal view moving in the direction opposite to that of the adapted stimulus, resulting in a repulsive perceptual aftereffect. However, this adaptation aftereffect disappeared when the adaptor was presented upside down (**Figure 2b**, mean difference = 0.010°, *t* (15) = 0.053, *p* = 0.958, Cohen’s *d* = 0.013, CI = [-0.388, 0.408], BF_10_ = 0.256). A two-way repeated-measures ANOVA was performed on the point of subjective equality (PSE), with the adapting direction (right +30°vs. left - 30°) and orientation of the BMs (upright vs. inverted) as within-subject factors, showing a significant interaction (*F* (1, 15) = 8.227, *p* = 0.012, *η*_p_^2^ = 0.354, BF_10_ = 15.721).

**Figure 2.**
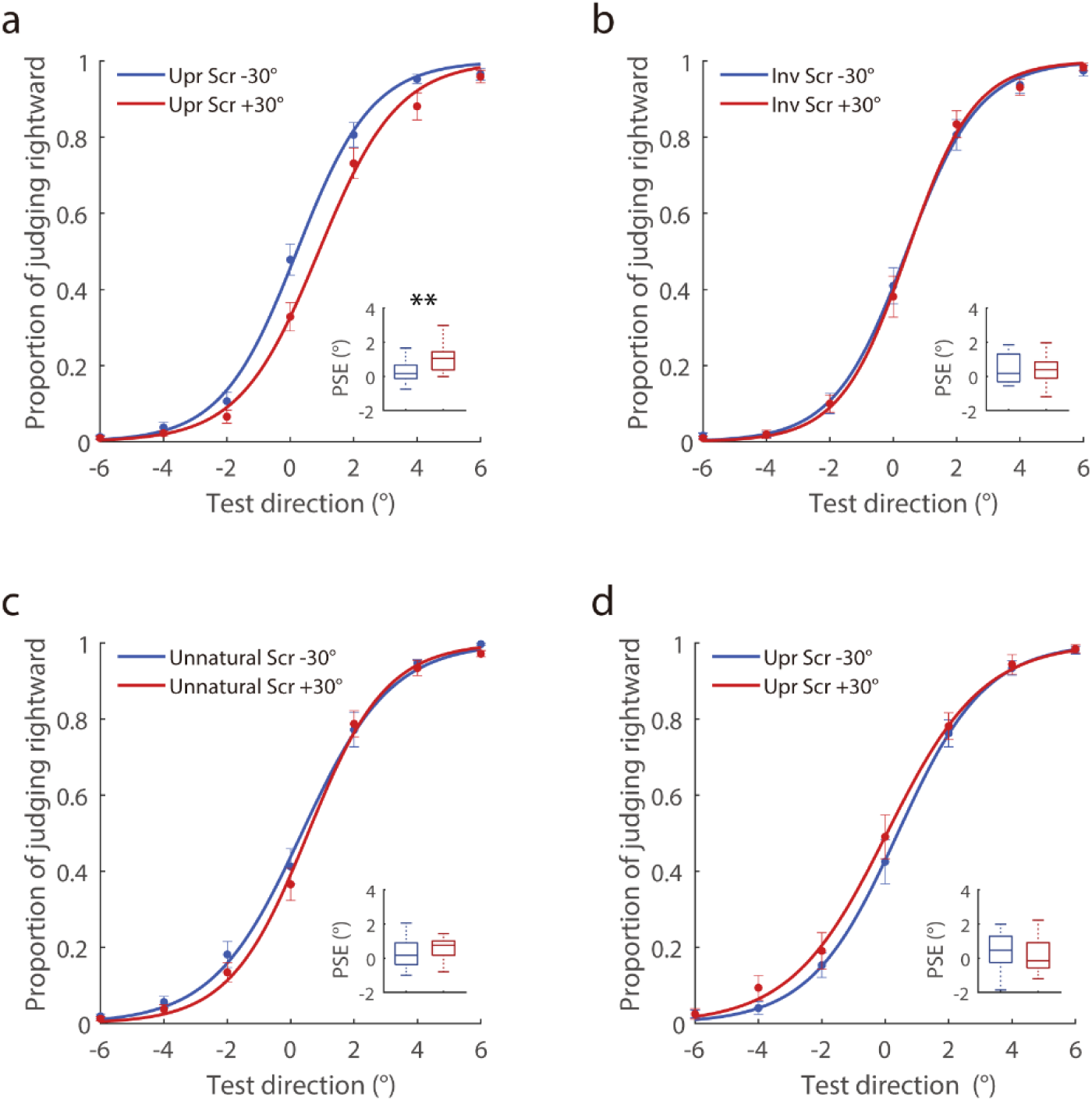
Results of Experiment 1. (**a-d**) The results of Experiment 1a-d are shown in four panels, respectively. The proportion of responses in which participants judged the rightward walking direction was plotted as a function of the presented walking direction. The red lines correspond to the rightward adapting direction (+30°), and the blue lines correspond to the leftward adapting direction (-30°). The inset boxplots depict the distribution of the corresponding points of subjective equality (PSEs) for each adapting direction. The central boxes represent the interquartile range (25th–75th percentiles), and the line inside the boxes represents the median. The upper and lower whiskers display the maximum and minimum data values, respectively. Asterisks indicate significant differences between the two adapting directions (\*\**p* < 0.01). Scr indicates the scrambled BM sequences used as adaptors.

Previous studies suggest that the gravitational acceleration pattern is critical for identifying life motion (Chang & Troje, 2009a; Troje & Westhoff, 2006). To verify this, we ran an additional control group in which the natural biological dynamics of the adaptors was disrupted by removing dots’ accelerations along the trajectory fragments (Experiment 1c). As expected, the adaptation aftereffect vanished (**Figure 2c**, mean difference = 0.243°, *t* (15) = 1.097, *р* = 0.290, Cohen’s *d* = 0.274, CI = [-0.229, 0.715], BF_10_ = 0.428). To further confirm that the observed walking direction aftereffect is specialized for life motion, we investigated whether adaptation generalizes from BM to non-biological object motion. A new group of participants adapted to the upright scrambled PLDs of a human walker and then judged the perceived moving directions of a rolling ball (Experiment 1d). There was no significant adaptation aftereffect (**Figure 2d**, mean difference = -0.355°, *t* (15) = -1.765, *р* = 0.098, Cohen’s *d* = -0.441, CI = [-0.784, 0.074], BF_10_ = 0.904). Taken together, our study supports the existence of neural representations that encode specific directions conveyed by local BM. Crucially, this mechanism is selective for biological kinematic constraints (e.g., gravity-consistent accelerations) and poorly transfers to non-biological motion.

Notably, a recent study using a similar visual adaptation paradigm with different BM stimuli did not find a perceptual aftereffect following adaptation to a scrambled point-light walker (Chen et al., 2023). As the directional information from local BM is inherently weaker than that from intact human walkers, the absence of an adaptation aftereffect in Chen’s study might be attributed to their small sample size (n = 8), which likely limited the statistical power to detect relatively subtle effects. To address this and verify the generality of our findings, we ran two supplemental experiments with 16 participants each (see details in **Supplemental Material**). In Supplemental Experiment 1, we scrambled the BM sequences both in spatial position and temporal phase during adaptation (i.e., each dot of the originally spatially scrambled point-light walker started at a random frame). In Supplemental Experiment 2, we adopted the BM sequences identical to those used in Chen’s study. As in our main experiments, a colour detection task was imposed during the adaptation period to ensure effective visual exposure. In line with our main findings, both supplemental experiments revealed significant adaptation aftereffects (**Figure S1**, Supplemental Experiment 1: mean difference = 0.409°, *t* (15) = 2.396, *р* = 0.030, Cohen’s *d* = 0.599, CI = [0.045, 0.773], BF_10_ = 2.240; Supplemental Experiment 2: mean difference = 0.290°, *t* (15) = 2.280, *р* = 0.038, Cohen’s *d* = 0.570, CI = [0.019, 0.561], BF_10_ = 1.875), which supports the robustness of direction adaptation driven by local biological kinematics, independent of specific stimulus configurations.

### Experiment 2: Cross-species adaptation to local BM

In Experiment 2, we investigated whether the observed adaptation aftereffect can occur in a cross-species manner. We had three groups of participants adapted to scrambled BM sequences of different terrestrial vertebrates (i.e., pigeon, cat and dog), respectively, and then evaluated their walking direction judgments of intact human walkers. Although the adaptors and test stimuli exhibited great differences in appearance, the PSEs in the rightward adapting condition were significantly higher than those in the leftward adapting condition across all three groups (**Figure 3a**), indicating cross-species adaptation aftereffects. A two-way mixed ANOVA comparing PSEs between the two adapting directions across three adaptation categories showed a significant main effect of adapting direction (*F* (1,45) = 24.902, *p* < 0.001, *η*_p_^2^ = 0.356, BF_10_ = 1712). The lack of a significant main effect of adaptation category (*F* (2, 45) = 0.832, *p* = 0.442, *η* ^2^ = 0.036, BF_10_ = 0.409) and the interaction effect (*F* (2, 45) = 0.977, *p* = 0.384, *η* ^2^ = 0.042, BF_10_ = 0.302) suggests a similar pattern of cross-species aftereffects observed across groups.

**Figure 3.**
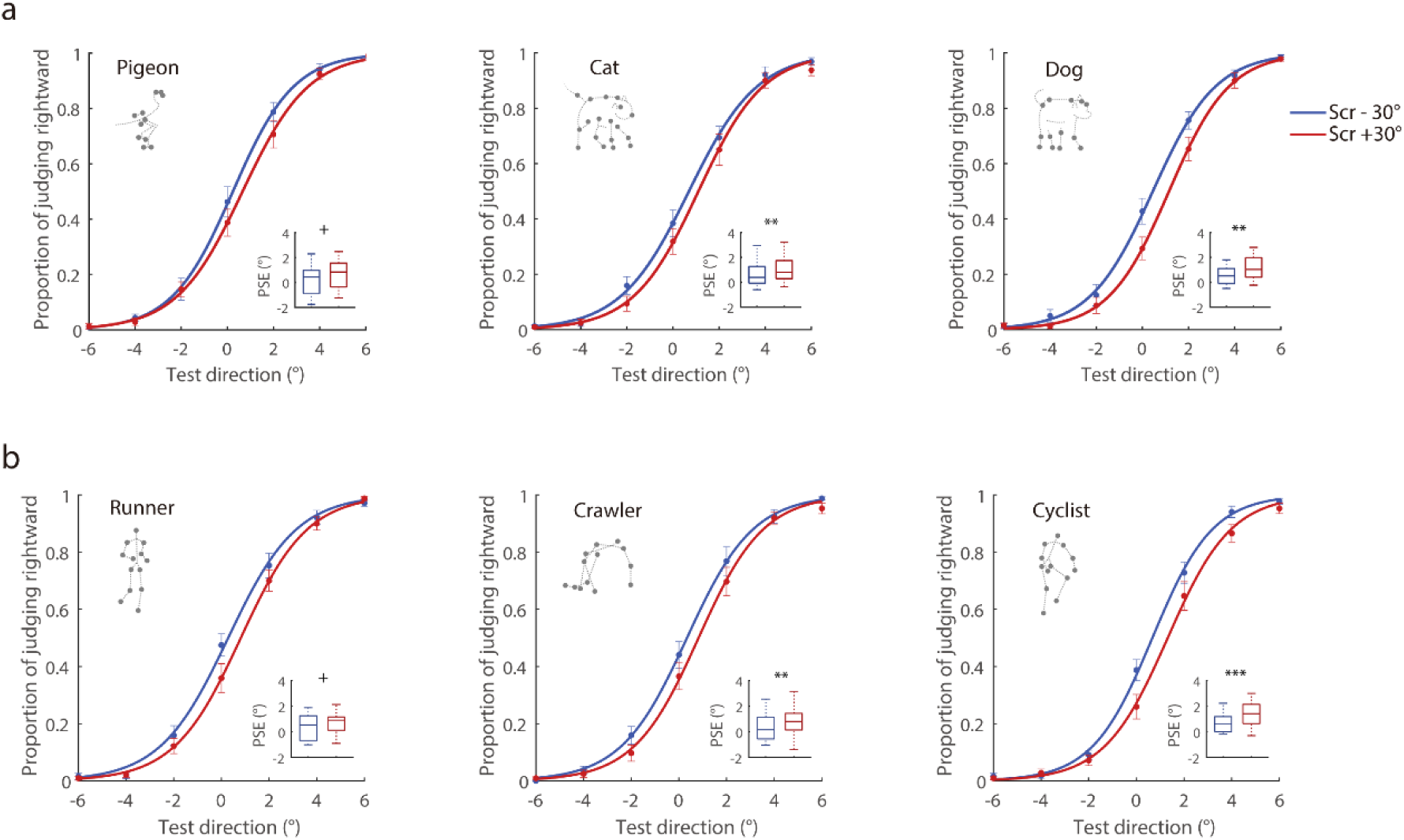
Results of Experiments 2 and 3. (**a**) Experiment 2: three groups of participants were adapted to different scrambled walking animals (pigeon, cat, dog) respectively, and tested with intact human walkers. (**b**) Experiment 3: another three groups of participants were adapted to different scrambled human actions (runner, crawler, cyclist) respectively, and tested with intact human walkers. Psychometric functions illustrate viewing direction judgments after adapting to various local BM sequences and inset boxplots present the distributions of the PSEs for each adapting direction. The top left corner of each panel shows an exemplar of the intact BM stimulus (oriented +30°) used to generate the spatially scrambled adaptors in each experimental group. Asterisks indicate significant differences between the two adapting directions (+*p* = 0.05, \**p* < 0.05, \*\**p* < 0.01, \*\*\**p* < 0.001). Scr indicates the scrambled BM sequences.

### Experiment 3: Cross-action adaptation to local BM

Experiment 3 tested whether the cross-category adaptation aftereffect would occur when the adapting and testing BM stimuli differed in action. Following a similar procedure as Experiment 2, another three groups of participants were tested for adaptation aftereffects while the adaptor actions were changed to running, crawling, and cycling, respectively. Despite noticeable kinematic differences among these actions (e.g., motion period, trajectories of key joints), there was a significant PSE difference between the two adapting directions for each group **(Figure 3b**), indicating cross-action adaptation aftereffects. A two-way mixed ANOVA was conducted with the adapting direction (±30°) as the within-subject factor and the adaptation category (runner, crawler, or cyclist) as the between-subject factor. Results showed a significant main effect of adapting direction (*F* (1, 45) = 29.312, *p* < 0.001, *η*_p_^2^ = 0.394, BF_10_ = 7100), but no significant main effect of adaptation category (*F* (2, 45) = 1.288, *p* = 0.286, *η*_p_^2^ = 0.054, BF_10_ = 0.550) or interaction effect (*F* (2, 45) = 0.553, *p* = 0.579, *η*_p_^2^ = 0.024, BF_10_ = 0.214). Together with previous findings, these results suggest the existence of a general neural mechanism representing specific moving directions of local BM in the human visual system, which is robust to changes in species and action.

### Summary of the adaptation aftereffects across all experiments

To further compare the strength of the aftereffects among experiments, we calculated the PSE shift for each experimental condition by subtracting the leftward-adapting PSE from the rightward-adapting PSE (a positive value indicates a repulsive aftereffect). **Figure 4** summarizes the mean PSE shifts, individual data distributions, and group estimates derived from 1,000 bootstrap samples of the original data (Davison & Hinkley, 1997). This visualization enables intuitive assessment of individual variability as well as robust estimation of group-level effect sizes across experimental conditions. Complete statistical results are detailed in **Table S1**. These findings unveiled robust adaptation aftereffects in all groups exposed to scrambled displays of natural BM from various animal and human movements. Conversely, such adaptation aftereffects were absent when the biological characteristics of the motion signals were disrupted (e.g., inverted and unnatural conditions), or when tested with object motion (e.g., test ball condition). This pattern underscores the specificity of directional adaptation effects for motion signals from living organisms across diverse species and action types.

**Figure 4.**
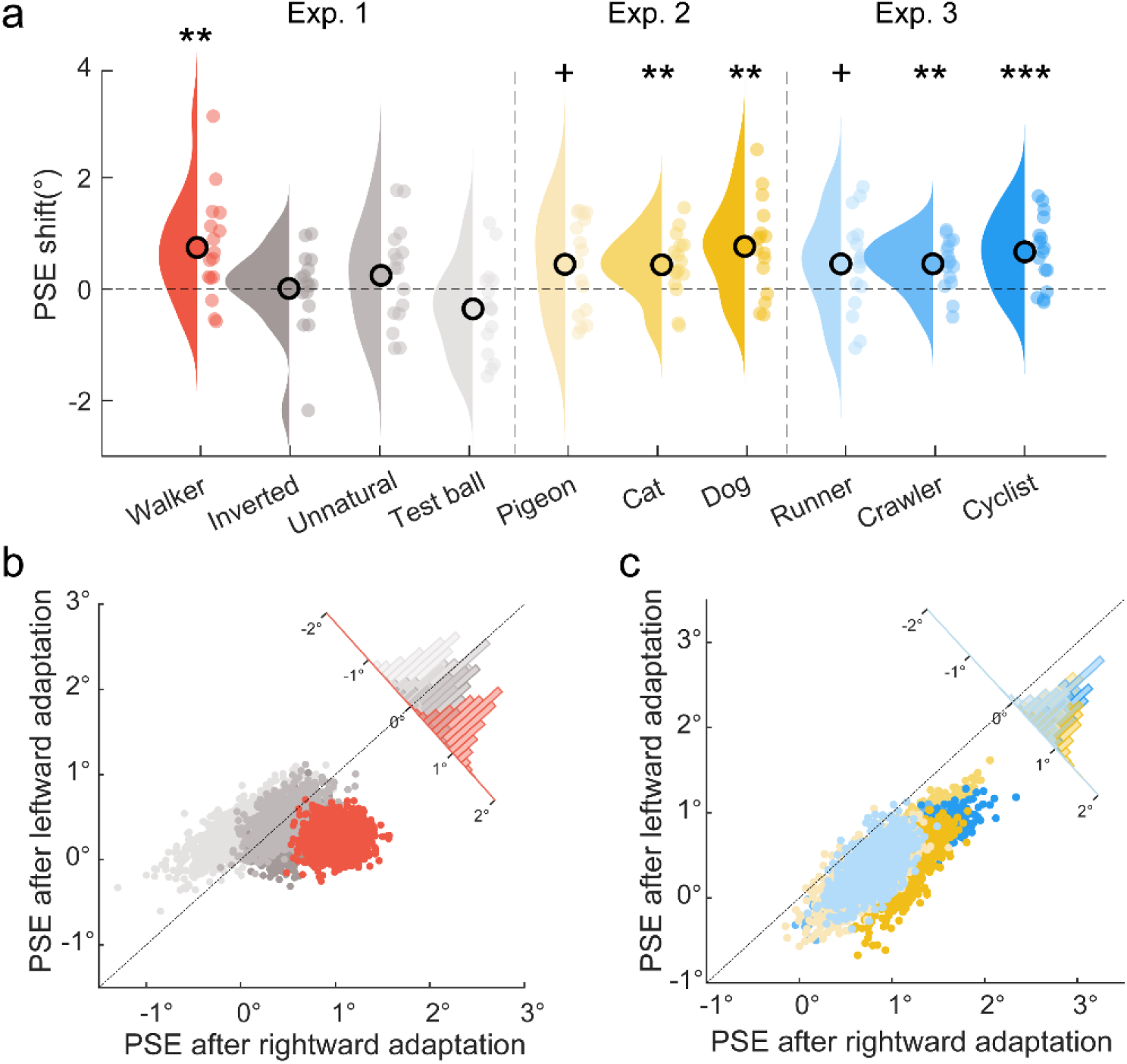
Summary of the adaptation aftereffects across experiments. (**a**) The PSE shifts are displayed separately for each of the ten experimental conditions. Individual data distributions (shaded region) were plotted for each group using the Raincloud plots package (Allen et al., 2021). Small coloured circles represent individual data points, and large open circles represent the means of PSE shifts. Asterisks indicate significant adaptation aftereffects (+*p* = 0.05, \**p* < 0.05, \*\**p* < 0.01, \*\*\**p* < 0.001). Bootstrapped PSEs in the leftward and rightward adaptation conditions are shown for (**b**) Experiment 1 and (**c**) Experiments 2-3. Bootstrapped samples falling below the diagonal line (e.g., coloured dots) indicate a repulsive adaptation aftereffect, whereas those proximal to the diagonal line (e.g., grayscale dots) suggest a null aftereffect. The histograms in the corner illustrate the distributions of the PSE shifts between the two adapting directions.

### Disentangling perceptual and decisional mechanisms in local BM direction adaptation through drift-diffusion modeling

Although we found robust and consistent PSE shifts across various local BM adaptation groups, it remains unclear whether these effects reflect genuine perceptual changes or merely decision-level biases. To address this issue, we applied the drift-diffusion model (DDM) to our behavioral data (see **Supplemental Material** for details). In the DDM framework, perceptual decision-making is described as the accumulation of noisy sensory evidence over time until a decision threshold is reached. By fitting participants’ choices and reaction times, DDM allows us to disentangle changes in perceptual processing—quantified by the drift rate (*v*, representing the efficiency of sensory evidence accumulation)—from decision-level factors such as prior response preference (starting point *z*) or response caution (decision bound *a*) (Ratcliff et al., 2016; Zhan et al., 2025). This quantitative dissociation provides a principled basis for interpreting adaptation aftereffects, offering unique insights into the underlying cognitive mechanisms (Witthoft et al., 2018). Our DDM analysis revealed significant adaptation effects on drift rate (*v*) across local BM conditions (**Figure 5a-c**; see detailed statistics in **Table S2**). Specifically, exposure to a particular motion direction reduced the efficiency of sensory evidence accumulation toward the adapted direction, thereby biasing perception away from the adaptor. This pattern aligns with the fatigue model of perceptual adaptation, which posits a suppression of sensitivity in neurons tuned to the adapted feature. Importantly, individual shifts in drift rate induced by adaptation were strongly correlated with PSE shifts (**Figure 5d**, *p* < 0.001), suggesting that the efficiency of sensory evidence accumulation plays a critical role in determining the strength of the behavioral aftereffect. In contrast, shifts in starting point (*z*) and decision bound (*a*) were minimal and did not reach statistical significance across experimental conditions (see **Figure S2** and **Table S2**). Furthermore, the repulsive effects on drift rate were specific to BM stimuli across species and actions, whereas control conditions (inverted, unnatural, and non-biological motion) showed no such effects. Together, these computational results provide compelling evidence that local BM adaptation is primarily driven by alterations in perceptual processing (i.e., sensory evidence accumulation) rather than shifts in decision criteria. This dissociation indicates that the observed behavioral aftereffects arise from a genuine recalibration of the visual system’s sensitivity to local biological kinematics.

**Figure 5.**
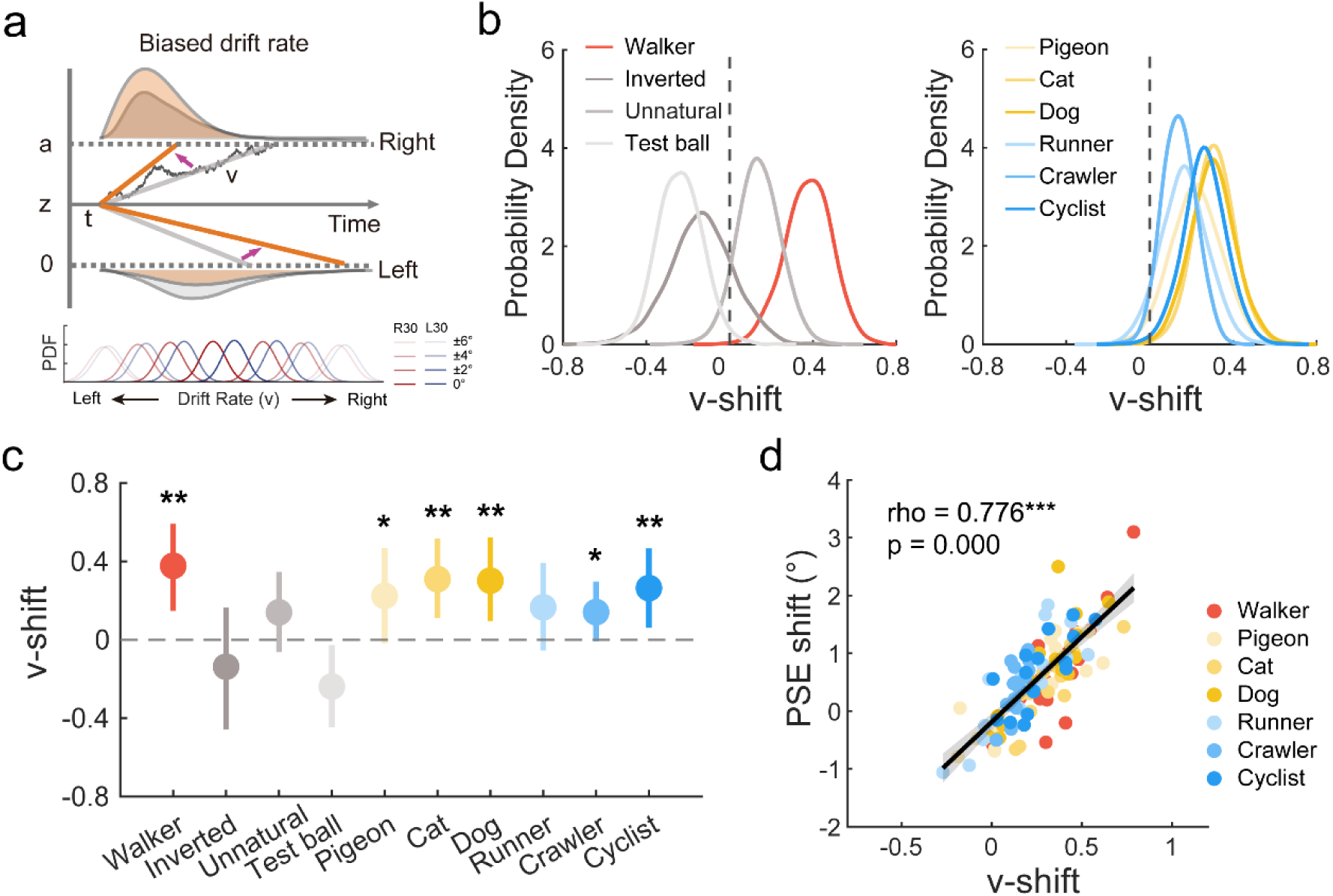
Drift-diffusion modeling results. (**a**) Schematic illustration of adaptation effects on the drift rate parameter (*v*) within the DDM framework. Perceptual bias is captured by alterations in the drift rate, reflecting the efficiency of sensory evidence accumulation toward different decision thresholds. The inset probability density functions (PDFs) depict the modulation of drift rates by leftward (L30, blue) and rightward (R30, red) motion adaptations for test stimuli at angular deviations of 0°, ±2°, ±4°, and ±6°. (**b**) Posterior distributions of the adaptation effect on drift rate for the best-fitting model in Experiments 1-3. The curves quantify this effect as the parameter differences between -30°and +30°adapting conditions across test directions, defined as *v*-shift = *v*_-30°_– *v*_+30°_. A significant repulsive effect is identified where the majority of the posterior mass (> 95%) falls above zero, indicating that the drift rate for the leftward adaptation condition is higher than that for the rightward adaptation condition. (**c**) Comparisons of adaptation-induced drift rate shifts across experiments. Error bars represent the 95% highest density interval of the parameter distribution. \**P* > 0.95, \*\**P* > 0.99. (**d**) Subject-level correlation between drift rate shifts and PSE shifts. Individual data points are overlaid with their linear regression lines and 95% confidence intervals. Spearman’s ρ effect size and p-value are reported. \*\*\**p* < 0.001.

### Correlations between perception of various local BMs

Lastly, we assessed individual abilities to process local BM cues across species and actions. Sixty-one participants performed a direction discrimination task with spatially scrambled displays of seven BM stimuli (human walker, pigeon, cat, dog, runner, crawler, and cyclist), each briefly presented in sagittal view (i.e., facing rightward or leftward). Results demonstrated that participants can readily retrieve information about heading direction from spatially scrambled BM sequences spanning diverse species and actions. Direction discrimination performance was significantly above chance for all local BM conditions (Wilcoxon signed-rank test, all *p* values < 0.001), with mean accuracies (mean ± SEM) of 80.3±2.0% (human walker), 97.0±0.7% (pigeon), 86.1±1.7% (cat), 91.1±1.2% (dog), 76.8±1.7% (runner), 92.1±1.2% (crawler), and 83.4±1.7% (cyclist), respectively. Furthermore, we also observed considerable individual variability in the ability to detect life motion. Spearman’s correlation analysis revealed significant pairwise correlations in direction judgment accuracy among different species and actions (**Figure 6a**, **6b**, all *p* values < 0.001), suggesting that the perceptions of diverse local BM cues are intrinsically linked at the individual level.

**Figure 6.**
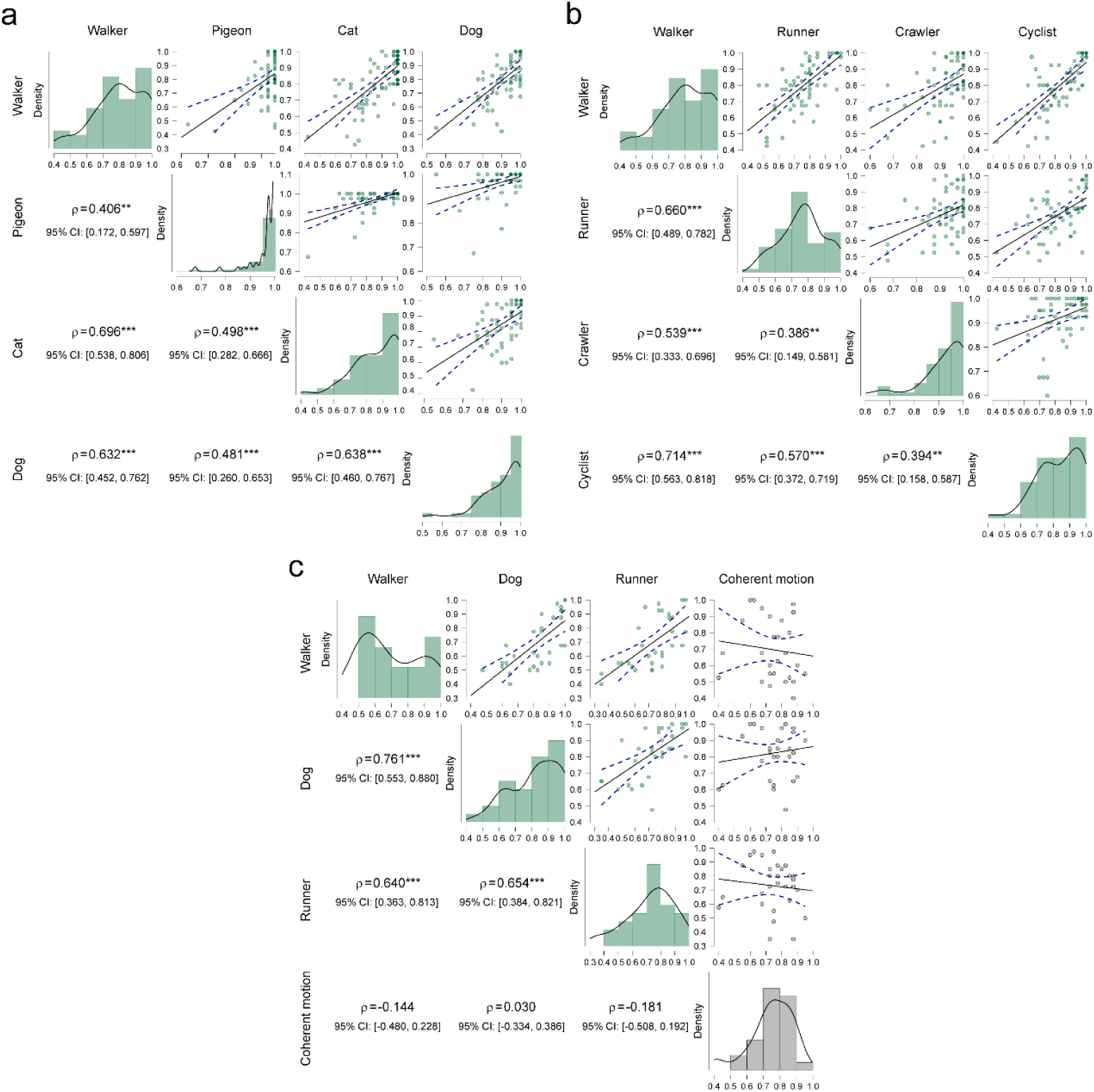
Correlations of direction discrimination performance on local BM across species and actions. **(a, b)** Correlation matrices for seven local BM stimuli tested in the main experiment, grouped by species categories and action categories, respectively. (**c**) Correlation matrix for the control experiment, comparing three representative local BM stimuli and one non-biological coherent motion stimulus. Individual data points in the upper triangle of the matrix are overlaid with their respective linear regression lines and 95% confidence intervals. The lower triangle of the matrix shows the corresponding Spearman’s *ρ* and 95% confidence intervals. Asterisks indicate significant correlations (\*\**p* < 0.01, \*\*\**p* < 0.001). The histograms along the diagonal illustrate the performance distribution of the corresponding BM category.

To determine whether these correlations were specific to local BM processing or reflected more general perceptual abilities, we conducted a control experiment in which a new group of thirty participants performed a similar direction discrimination task on two distinct types of stimuli: local BM from three representative categories (scrambled displays of human walking, running, and dog locomotion), and non-biological coherent motion (random-dot kinematogram) as a critical baseline control. Direction discrimination performance was significantly above chance for all conditions (Wilcoxon signed-rank test, all *p* values < 0.001), with mean accuracies (mean ±SEM) of 69.6±3.4% (human walker), 73.0±3.1% (runner), 82.3±2.7% (dog), and 74.9±2.4% (coherent motion) respectively. Results showed significant positive correlations among all pairs of local BM stimuli (**Figure 6c**, all *p* values < 0.001), replicating the key findings observed in the primary experiment. In contrast, no significant correlations were found between any local BM condition and the non-biological coherent motion condition (all *p* values > 0.3), suggesting that the observed associations are specific to BM processing rather than attributable to general motion perception, attention, or task engagement. Taken together, these correlation patterns corroborate the cross-category adaptation results, supporting a shared neural mechanism underlying the perception of life motion signals in terrestrial vertebrates.

## Discussion

The debate in the field of BM perception regarding the contributions of local biological kinematic information versus global motion-mediated configural information has been ongoing. Recently, accumulating evidence has suggested that local kinematic cues provide key clues (e.g., animacy and moving direction) for BM perception and could be processed independently of global configuration (Chang & Troje, 2009a; Troje & Westhoff, 2006). It is proposed that there might exist a unique visual mechanism sensitive to local BM cues, enabling organisms to effectively detect and respond to living entities in their visual environment. Thus, it is reasonable to hypothesize that the directional information carried by local kinematic cues, as a highly salient feature for life detection, may be selectively processed in the visual system. The current study employed the visual adaptation paradigm in a series of experiments to examine the neural direction encoding in response to local biological kinematics, testing its specificity and invariance, thereby advancing our understanding of the life motion detection.

First, we found that adaptation to a scrambled point-light walker significantly biased the perceived direction of subsequently presented human walker. Further mechanistic insights from drift-diffusion modeling indicated that this perceptual bias was rooted in alterations to sensory evidence accumulation (quantified by changes in drift rate) rather than shifts in decision criteria. Notably, individual variations in drift rate reliably predicted the magnitude of the observed perceptual aftereffects. These findings provide direct empirical evidence supporting the existence of neural mechanisms that are sensitive to direction information derived from local BM cues. What specific visual cue is exploited to elicit locomotion direction perception? The movements of terrestrial animals are inevitably constrained by gravity, resulting in local velocity and acceleration profiles jointly shaped by biomechanical and gravitational forces. These distinctive gravity-dependent dynamics, especially arising from feet motion, provide vital cues for detecting living organisms (Troje & Chang, 2023). Consistent with this framework, our control experiments revealed that the aftereffect disappeared once these gravity-compatible kinematic signatures were disrupted (e.g., by vertically flipping the scrambled PLDs or modifying joint trajectories to constant motion). Moreover, the absence of the cross-category adaptation aftereffect between local BM and non-BM cues (i.e., rolling ball) aligns with previous findings that motion direction adaptation does not generalize between human walkers and rotating spheres (Chen et al., 2023), suggesting that the observed perceptual aftereffect is unlikely to be accounted for by a generic representation of object-level moving direction. Hence, converging evidence highlights the existence of direction-sensitive representations in the human brain that are specifically tuned to biological characteristics embedded in life motion.

From an evolutionary perspective, a quick visual alert system tuned to life motion for identifying animate agents is crucial for species’ survival and everyday interactions, such as seeking caregivers and avoiding predators. Extensive work has shown that human sensitivity to local BM is not restricted to conspecific motion. For instance, human can readily retrieve direction information from scrambled displays of biped and quadruped animals (Chang & Troje, 2008). Newly hatched chicks exhibit a spontaneous preference toward BM patterns, which even extends to potential predators such as cats (Vallortigara et al., 2005). This cross-species sensitivity suggests that locomotion direction may be represented in a category-general manner, a mechanism likely shaped by long-term exposure to diverse BM patterns in a gravitational environment through evolution. The current study demonstrated that direction aftereffects could manifest during cross-category adaptation between different species or various actions, despite considerable dissimilarities in physical characteristics (e.g., the number of lower limbs, amplitude of motion, gait frequencies etc). We further showed that individual abilities to discriminate heading direction across diverse local BM patterns are intrinsically linked, implying a shared, invariant neural representation for local BM in the human visual system. It is worth noting that, in contrast to action-selective tuning for global BM processing revealed by adaptation to intact point-light animations (Van Boxtel & Lu, 2013), the action-invariant coding observed for local kinematic cues indicates differentiated processing of local BM signals.

Direction information retrieved from local BM signals should not be simply categorized as a low-level physical attribute; rather, it reflects high-level processing related to social functions. In a social world, heading direction provides important clues for assessing others’ goals and reacting appropriately. A growing body of research has shown that local BM cues can act as a salient feature for the visual system and exert a robust reflexive attentional orienting effect even without subjective awareness of the biological nature (Wang et al., 2010, 2014). In addition, local motion cues may function as an effective signal for the presence of living creatures in the environment. Chang et al. (2008) have found that the ability to discriminate local motion direction is correlated with perceived animacy from scrambled displays. Furthermore, sensitivity to local BM signals has been considered a potential hallmark of social cognitive skills. A recent twin study has revealed that the ability to identify local BM cues is highly heritable and correlated with individual autistic traits (Wang et al., 2018).

Perception of point-light animations can invoke a broad cortical network along the motion and form pathways (e.g., middle temporal complex, fusiform gyrus), and then extend to higher-level cortices specialized for socially relevant information processing (e.g., superior temporal sulcus (STS), temporoparietal junction and inferior frontal gyrus) (Pitcher & Ungerleider, 2021). As a central hub of the social brain network, STS plays a critical role in processing dynamic social cues, such as facial expressions, eye gaze, and body movements (Yokoyama et al., 2021). Specifically, Jellema et al. (2004) found neurons in the macaque STS sensitive to biological walking directions. Human neuroimaging work has demonstrated an orientation-dependent representation of BM in the STS (Grossman et al., 2000), and this region also responds to scrambled BM (Jastorff & Orban, 2009). In light of a number of developmental studies showing that sensitivity to BM emerges early in life, the subcortical network, regarded as evolutionarily conserved and early maturing, is presumed to be engaged in life motion detection (Troje, 2008). Recent animal work has revealed subcortical responses for animacy cues on newly hatched chicks without any prior visual experience (Lorenzi et al., 2024). A human neuroimaging study has discovered that a subcortical thalamic area is sensitive to local kinematic information and can discriminate between intact and perturbed local kinematics (Chang et al., 2018). Recent fMRI work on humans and macaques has shown that the superior colliculus exhibits selective responses to local biological kinematics (Lu et al., 2024).

Hirai and Senju have proposed a two-stage model for BM processing (Hirai & Senju, 2020). This model includes an initial “step detector” stage, which rapidly processes local BM cues and is located in the subcortical network, and a later “bodily action evaluator” stage, which slowly processes the fine global structure-from-motion information and largely relies on cortical circuits. Compatible with this view, our results point to a unique mechanism for processing local motion cues that operates independently of global configuration. We speculate a hierarchical process: direction information is initially extracted by a rapid visual filter which is sensitive to some crude perceptual features shared in diverse life motion signals (e.g., the gravity-dependent dynamics) for animacy detection, and then fine-tuned by integrating global configurational cues to identify body actions for more complex social cognition. Specifically, a subcortical-cortical brain network might be involved in this hierarchical process. Future work may systematically characterize the detailed tuning properties of local BM processing across a full range of directions, and employ neuroimaging approaches to reveal how local and global BM cues are represented and potentially interact within the neural network.

In summary, using various types of local BM cues, we have discovered robust moving-direction adaptation aftereffects specialized for life motion signals. Our results indicate the existence of a neural mechanism representing specific heading directions of local BM in human visual system, and this representation exhibits invariance across different species and action types, supporting and enriching the “life motion detector” theory. These findings not only provide novel insights into the neural mechanisms underlying evolutionary-adaptive coding of BM signals, but also have potential practical implications for the development of algorithms detecting and imitating life motion effectively.

## Methods

### Participants

A total of 181 young adults participated in the adaptation experiments. Five observers were excluded from the analysis due to poor performance in discriminating the walking direction of the intact point-light walker (goodness of fit, R^2^ value below a cutoff of 0.85). Experiments 1-3 included 144 participants (83 females, mean age = 23.2±2.7 years), who were randomly assigned to the three primary adaptation experiments, each comprising three groups of 16 participants. Additionally, 32 individuals (16 females, mean age = 23.3±1.8 years) engaged in the two supplemental adaptation experiments, with each experiment consisting of 16 participants. Sample size was determined based on a priori power analysis (matched-pairs *t* test, two-tailed) using G*Power (Faul et al., 2009). Given the lack of prior studies matching our experimental designs, we chose a moderate effect size (Cohen’s *d* = 0.75), α = 0.05, and 80% power, informed by our pilot data, which yielded a recommended sample size of 16 per group.

A total of 91 participants (40 females, mean age = 23.8±2.8 years) took part in the local BM perception test. 61 individuals from the adaptation experiments completed an additional session to assess their direction discrimination abilities for various local BM patterns. 30 naïve participants performed the local BM test alongside a non-biological coherent motion discrimination task, and this experiment was preregistered on the Open Science Framework (https://doi.org/10.17605/OSF.IO/UK3YE) prior to data collection.

All participants were naive to the purpose of the study, had normal or corrected-to-normal vision and provided written informed consent. This study was conducted in accordance with the Declaration of Helsinki and approved by the institutional review board of the Institute of Psychology, Chinese Academy of Sciences (Protocol Number: H21058).

### Stimuli and Apparatus

In the adaptation experiments, the test stimuli were intact point-light displays (PLDs) of a human walker (all experiments except Experiment 1d) or a rolling ball (Experiment 1d). The human walker, adopted from Troje (2002), was depicted by a set of 13 dots representing the head and critical joints (shoulders, elbows, wrists, hips, knees and ankles). This display subtended 6.8°×3.7°in visual angle (sagittal view) and contained 30 frames in each 1-s gait cycle. To avoid participant prediction, the initial frame of the PLDs was randomized for each trial. The rolling ball was composed of 150 dots, had a diameter of 5.5°, and rolled along a virtual horizontal axis at an angular velocity of 60° per second, similar to that of the BM walker (Cheng et al., 2022). Each dot was 0.2°in diameter. The position of the testing stimulus was jittered by 0°to 1.3°from the center of the screen on each trial. The test stimuli, either human walkers or rolling balls, were projected onto a 2D plane. Although this 2D projection may result in bistable perception (moving toward or away) due to the lack of depth cues, we implemented specific measures to minimize such ambiguity and argued that individual differences in perceived direction do not affect the validity or interpretation of adaptation effects (see **Procedure** for details).

Scrambled BM sequences, derived from intact sequences, served as adaptors in the adaptation experiments or as test stimuli in the local BM perception task. For every scrambled BM stimulus, we randomly assigned the starting position of each dot within a spatial envelope that was empirically determined by visual matching for each category. This envelope was set to about 1.2 times the height and 1.4 times the width of the bounding box of the test stimulus (i.e., the intact human walker), ensuring adequate spatial coverage of the test stimulus by the adaptors. To keep the full motion trajectory of each dot within the envelope, the range of possible starting positions was adjusted by subtracting the dot’s maximal movement displacement. For each trial, a new scrambled pattern was generated with independent randomization of dot positions. This scrambling procedure disrupted the global form of the BM stimuli while preserving the local motion signals, following established methods in previous studies (Troje & Westhoff, 2006; Wang et al., 2018). Scrambled PLDs of a human walker were used in Experiment 1. The inverted scrambled stimulus was obtained by mirror flipping the upright scrambled PLD vertically. We also introduced unnatural local motion, which was derived from the fragments identical to the scrambled sequences but with the key biological characteristic of dynamics (i.e., motion acceleration) disrupted. That is, each dot moved along the trajectory path at a constant speed equal to the average speed of the corresponding dot.

Scrambled PLDs of three different animals, including a walking pigeon, cat (Troje & Westhoff, 2006) and dog, were employed in Experiment 2. The PLDs of the pigeon, cat, and dog were composed of 11, 18, and 14 markers, respectively. The gait frequencies were 1 Hz for the pigeon, 1 Hz for the cat, and 0.97 Hz for the dog. The walking dog BM stimuli were generated using a motion capture and tracking system (Nokov, Mars 4H). We obtained the PLD sequence of a German shepherd at a sampling rate of 60 Hz. Fifteen motion capture markers were attached to the dog’s anatomy: one each on the ear and nose, two on the back, two on the tail and two on each limb. The dog was allowed to walk freely within a confined space to capture its natural walking pattern. The three-dimensional coordinates of these markers were then extracted from the system. Following a smoothing process, these refined coordinates (31 frames per cycle) served as the dog BM stimulus. We excluded one marker located at the tail that contained irrelevant emotional motion, resulting in a final stimulus of 14 dots used in the present study. The full PLDs (sagittal view) approximately subtended visual angles of 6.5°×4.8°, 3.3°×7.0°, and 2.5°×4.6°for the pigeon, cat, and dog, respectively.

Scrambled PLDs of three different actions, including running (Chang & Troje, 2009a), crawling and cycling (Vanrie & Verfaillie, 2004), were used in Experiment 3. Each action consisted of 13 markers. The moving frequencies were 1 Hz, 0.63 Hz and 0.86 Hz, and the visual angles of the full PLDs were approximately 7.4°×3.0°, 3.5°× 7.4°, and 7.5°×3.7°for the runner, crawler, and cyclist, respectively. We selected these actions because they are axisymmetric and have clearly definable moving directions.

In the coherent motion discrimination task, participants viewed a cloud of 100 small dots (each subtending 0.1°of visual angle) presented randomly within a circular aperture of 8° diameter. All dots travelled at a speed of 10°/s. On each trial, a certain proportion of dots (signal dots) moved consistently in one direction (either leftward or rightward), while the remaining dots moved in random directions. The proportion of signal dots relative to the total number of dots, termed motion coherence, was fixed at 20% in this task.

All stimuli were presented as white dots against a gray background on a 27-inch LCD monitor (2560 ×1440 resolution, 60 Hz refresh rate) at a viewing distance of 60 cm. Experiments were controlled by the Psychophysics Toolbox extensions (Brainard, 1997; Pelli, 1997) for MATLAB (The MathWorks, Natick, MA).

### Procedure

#### Adaptation experiments

Each adaptation session comprised eight blocks for two adaptation conditions (adaptation to the left or right direction). Each block started with a 20-s pre-adaptation during which a scrambled BM sequence heading toward a certain direction (−30° leftward or +30°rightward) was displayed (**Figure 1**). The colour of a small part of the scrambled PLD switched to yellow for 333 ms every 3-5 seconds. Participants were asked to carefully track the movement of the adapting stimulus and indicate the colour change as quickly as possible. This covert task helped participants maintain attention during the whole adaptation period. In the following trials, after 0.6-s fixation on a central cross (0.4°× 0.4°), a test stimulus of an intact human walker or a rolling ball was presented for 500 ms. The test stimuli were shown with seven different moving directions in separate trials (left −6°, −4°, −2°, front 0°, right +2°, +4°, +6°). Participants were required to discriminate whether the test stimulus was heading leftward or rightward, prioritizing accuracy while responding as quickly as possible. After responses and a 0.6-s intertrial interval, participants were exposed to a 4-s additional topping-up adaptation and performed the same colour detection task (20% trials during topping-up adaptation exhibited one colour change). To minimize low-level adaptation, the center of the adaptor was randomly relocated every gait cycle within a 1.3°× 1.3° area. Importantly, participants were not explicitly informed about the direction information conveyed by the local BM stimuli during the adaptation experiments, ensuring they remained unaware of this aspect of the stimulus. Each adaptation condition contained 20 repetitions for each of the seven testing directions, resulting in a total of 140 trials which were presented in pseudo-random order and distributed in four sequential blocks (35 trials per block). Participants rested for at least half a minute between blocks. The order of the two adaptation conditions (i.e., adapting to ±30° directions) was counterbalanced among participants. To allow the visual system to recover from the adaptation, there was a minimum of a 5-min rest break between adaptation conditions.

Prior to the formal experiments, observers completed a brief practice session to familiarize themselves with the testing stimuli and the procedure. This session consisted of a 30-trial direction discrimination task using upright intact human walkers (or rolling balls for Experiment 1d) with six supraliminal directions (±5°, ±10°, ±15°; 5 repetitions each, 1-s presentation per animation), followed by 35 adaptation practice trials using adaptors heading along the sagittal axis without lateral deviation. These practices ensured that participants were capable of performing these tasks, meeting the criterion that correct response rates for ±6°test walkers/balls exceeded 80% and colour detection accuracy reached 80%.

This practice session also helped ensure that each participant maintained a clear and stable perception of the assigned test stimulus as moving toward or away from themselves. The brief 500-ms presentation duration minimized perceptual switching during the experiments. Although the test stimulus could be perceived as moving toward or away from the observer, adaptation effects can still occur as long as the perceived depth direction is consistent within individuals during adaptation and test phases, regardless of the absolute depth direction. Notably, most participants (approximately 90%) naturally perceived the walker as approaching rather than receding due to a strong facing bias for social agents (Vanrie et al., 2004). Critically, robust adaptation effects were observed for the intact human walker across all conditions, whereas the rolling ball control condition yielded no significant repulsive aftereffect even when restricted to participants who perceived the ball as approaching (n = 11). This collective evidence mitigates concerns that heterogeneity in depth perception confounded the results.

#### Local BM perception test

We conducted a separate perceptual test session using a within-subjects design to examine individual direction discrimination performance across different motion stimuli. After the adaptation experiments, 61 participants completed a direction discrimination task on a separate day, using spatially scrambled displays of seven BM sequences spanning different species and actions (human walker, pigeon, cat, dog, runner, crawler, and cyclist). The seven categories of scrambled BM sequences were assigned to separate test blocks with randomized order across participants. Participants rested for at least 30 seconds between blocks. Each block contained 80 trials of the same stimulus category: 20 trials for each of four moving directions (± 30°or ± 90°), presented in a pseudorandom order. We primarily report the results for stimuli presented at ±90°(left/right directions), as this type of stimuli has been used to probe local BM processing in previous literature (Chang & Troje, 2008; Troje & Westhoff, 2006). Similar results were obtained for the side view stimuli (±30°). A separate group of 30 naïve participants also completed a direction discrimination task for four types of motion stimuli: scrambled human walker, scrambled dog, scrambled human runner, and coherent motion (random-dot kinematogram). Stimuli from the four categories were presented in separate blocks with randomized block order. Each block contained 40 trials (20 leftward and 20 rightward movements) in randomized presentation order.

For the perceptual tests, each trial began with a 500-ms fixation cross (0.4°×0.4°), followed by a central stimulus displayed for 1000 ms. Participants were asked to report the stimulus moving direction (left or right) by pressing the corresponding arrow key. All participants received instructions and completed practice trials to ensure task comprehension prior to formal testing. No feedback was provided throughout the experiments.

### Data analysis

#### Quantifying the adaptation aftereffect

We calculated the proportion of responses indicating a rightward moving direction for the test stimuli (an intact human walker or rolling ball). Individual participants’ data were fitted with a Boltzmann sigmoid function: f(x) = 1/(1 + exp (- a × (x - c))), where x denotes the actual moving direction and c represents the point of subjective equality (PSE). The PSE is the moving direction of the test stimuli where the probabilities of leftward and rightward judgments are equal. If the average PSE under the rightward adapting condition is significantly greater than that under the leftward adapting condition, it reflects a repulsive adaptation aftereffect. We defined the PSE shift by subtracting the leftward-adapting PSE from the rightward-adapting PSE. A positive value indicates a repulsive adaptation aftereffect. In all experiments, we assessed the goodness of fit (R^2^) of the fitted Boltzmann function within each adapting direction.

#### Statistical analysis

We used JASP (version 0.95.0.0) to perform repeated-measures ANOVA, Spearman’s correlation, and paired-samples *t* tests. Greenhouse-Geisser correction was applied for violations of the spherical assumption. Bayes factors (BF_10_) were reported to quantify evidence for the alternative hypothesis (H_1_) relative to the null hypothesis (H_0_). Specifically, BF_10_ > 1 suggests evidence for the alternative hypothesis, with BF_10_ > 3 considered substantial evidence. Conversely, BF_10_ < 1 provides evidence supporting the null hypothesis, with BF_10_ < 1/3 considered substantial evidence for the null.

## Acknowledgments

This work was supported by grants from the Brain Science and Brain-like Intelligence Technology – National Science and Technology Major Project (2021ZD0203800), the National Natural Science Foundation of China (32430043), the Youth Innovation Promotion Association of Chinese Academy of Sciences (2018115), the Key Research and Development Program of Guangdong, China (2023B0303010004), and the Fundamental Research Funds for the Central Universities.

