## Supplemental materials for "A Generalized Life-Motion Mechanism Supports Invariant Directional Coding of Local Biological Kinematics in Humans"

#### Supplemental Methods

##### Stimuli and procedures for supplemental experiments

*Supplemental Experiment 1*

The stimuli and procedure were identical to those in Experiment 1a except that the point-light walker adaptors were scrambled both spatially and temporally. Specifically, the initial position of each dot was randomized within the display area, and each dot started at a random frame within each animation (**Figure S1a**).

*Supplemental Experiment 2*

Point-light stimuli, consistent with Chen et al.'s study (2023) and obtained from the Carnegie Mellon University motion-capture database (http://mocap.cs.cmu.edu), depicted the locomotion of a human walker with a set of 41 dots. Raw motion capture data were converted to 3D coordinates using the Biomotion toolbox (van Boxtel & Lu, 2013). The gait frequency of biological motion (BM) was 0.58 Hz (104 frames per gait cycle). The adapting stimuli (**Figure S1b**) were spatially and temporally scrambled from the intact BM presented in a side view (±30°). The spatiotemporal scrambling manipulation was achieved by randomizing both the starting position and the initial frame of each dot within each animation. The height of the adaptor stimulus was approximately 10°, and the height of the test stimulus was 8°. Each dot was rendered in white with a diameter of 2.2 arcmin. All stimuli were presented in the middle of the screen against a black background on a 20-inch CRT monitor (1024 × 768 resolution; 60 Hz refresh rate) at a viewing distance of 60 cm.

Supplemental Experiment 2 followed the same procedure as in Experiment 1a, apart from the following modifications. The adapting and test stimuli were changed to a different set of point-light BM sequences (see details above). Due to the changes in stimulus appearance, minor modifications were made to the colour detection task to match the task difficulty. That is, during the adaptation period, the colour of 8 randomly chosen dots (20% of the BM markers) intermittently switched from white to green.

#### Hierarchical drift-diffusion model analysis

While adaptation aftereffects on perceptual judgments are well-documented, whether these effects arise from changes in sensory processing or decision processes remains actively debated (Storrs, 2015). We applied a drift-diffusion model (DDM) to investigate the computational mechanisms underlying direction adaptation effects in the speeded two-alternative forced-choice task. Specifically, we modeled participants' reaction times (RTs) and choices across all adaptation conditions in Experiments 1-3, aiming to dissociate perceptual and decision-related contributions to the observed aftereffects. In this framework, decisions about moving direction are conceptualized as the accumulation of noisy sensory evidence over time from an initial state toward one of two decision boundaries (left vs. right direction). The DDM framework includes four key parameters: drift rate (*v*, reflecting the speed of sensory evidence accumulation), starting point (*z*, indicating a prior response bias), decision bound (*a*, representing the amount of evidence required to make a choice), and non-decision time (*t*, accounting for sensory encoding and motor execution processes).

Model fitting was implemented in Python 3 using the hierarchical drift-diffusion model (HDDM) toolbox (Pan et al., 2025; Wiecki et al., 2013). Prior to modeling, trials with RTs greater than 3 seconds were excluded, resulting in the removal of 145 trials across all experiments (0.32% of total, mean excluded RT = 5.23 s). Stimulus coding was applied such that decision boundaries corresponded to leftward (0) and rightward (+1) responses. All model parameters were assigned default informative priors as implemented in HDDM, and the outlier probability was set to 5%. Model parameters were specified using HDDM's regressor approach to incorporate experimental factors. Using hierarchical Bayesian estimation, subject-level and group-level parameters were simultaneously estimated using Markov-Chain Monte-Carlo (MCMC) methods. MCMC sampling was conducted with 10,000 samples (burn-in = 5,000; thinning = 4; chains = 3).

To systematically investigate how adaptation and test stimulus influenced the perceptual decision process, we constructed and compared a nested set of candidate models. These models explored various hypotheses regarding how the key DDM parameters (drift rate *v*, starting point *z* and decision bound *a*) were either fixed or modulated by adaptation condition (±30° direction) and test stimulus (0°, ±2°, ±4°, ±6°). We focused on these three parameters because they map directly onto the theoretical distinction between perceptual ($v$) and decisional ($z$, $a$) mechanisms: $v$ indexes perceptual processing efficiency (rate of evidence accumulation), whereas $z$ and $a$ reflect initial decision bias and response caution, respectively. In total, we compared 12 candidate models in a nested framework comprising one baseline model and three model families, as detailed below.

Baseline Model

m1: $z \sim 1$, $a \sim 1$, $v \sim 1$

Decision bias model (modulating starting point $z$):

m2: $z \sim 1$ + adaptation

m3: $z \sim 1$ + adaptation + test

Decision criterion model (modulating decision bound $a$):

m4: $a \sim 1$ + adaptation

m5: $a \sim 1$ + adaptation + test

Decision strategy model (modulating both starting point $z$ and decision bound $a$):

m6: $z \sim 1$ + adaptation, $a \sim 1$ + adaptation

m7: $z \sim 1$ + adaptation + test, $a \sim 1$ + adaptation

m8: $z \sim 1$ + adaptation, $a \sim 1$ + adaptation + test

m9: $z \sim 1$ + adaptation + test, $a \sim 1$ + adaptation + test

Dual mechanism model (extending the three best-performing models from m1-m9 by additionally modulating drift rate $v$):

m10: $z \sim 1$ + adaptation + test, $v \sim 1$ + adaptation + test

m11: $z \sim 1$ + adaptation + test, $a \sim 1$ + adaptation, $v \sim 1$ + adaptation + test

m12: $z \sim 1$ + adaptation + test, $a \sim 1$ + adaptation + test, $v \sim 1$ + adaptation + test

Convergence of the MCMC sampling was evaluated with the Gelman-Rubin R-hat statistic. Model fits were evaluated using WAIC and LOO-CV which showed perfectly consistent rankings (Spearman's rho = 1.0, $p < 0.001$). Thus, only WAIC results are reported below (**Figure S3a**). Model comparison revealed that including drift rate *v* as a modulated parameter substantially improved model performance, suggesting that adaptation and test stimuli primarily modulated the rate of evidence accumulation. Model m11 ($z$ ~ 1 + adaptation + test, $a$ ~ 1 + adaptation, $v$ ~ 1 + adaptation + test) showed the best fit, with successful convergence (all R-hat < 1.05 for Experiments 1-3) and the lowest WAIC deviance (WAIC = 166.05), demonstrating clear superiority over competing models ($\Delta$WAIC > 14.88). Posterior predictive checks further validated model adequacy by demonstrating strong correspondence between simulated and observed choices and RT distributions. The simulated RTs and choices, generated from subject-level parameters of the best-fitting model and matching the number of trials and sample sizes in the behavioral experiment, effectively captured performance across various testing stimulus sets and adaptation conditions (**Figure S3b**).

For our primary analyses, we focused on adaptation-induced changes, with testing direction included as a covariate in the model. We examined how adaptation influenced the key DDM parameters ($v$, $z$, $a$). Adaptation effects were quantified as the difference between the parameter values estimated under the leftward-adapting (-30°) condition and those under the rightward-adapting (+30°) condition (i.e., *Parameter*-shift = $Parameter_{-30°} - Parameter_{+30°}$). Positive values for drift rate ($v$) and starting point ($z$) indicate repulsive aftereffects (i.e., a biased process away from the adaptor direction), whereas changes in decision bound ($a$) reflect shifts in response caution.

### Supplemental Figures

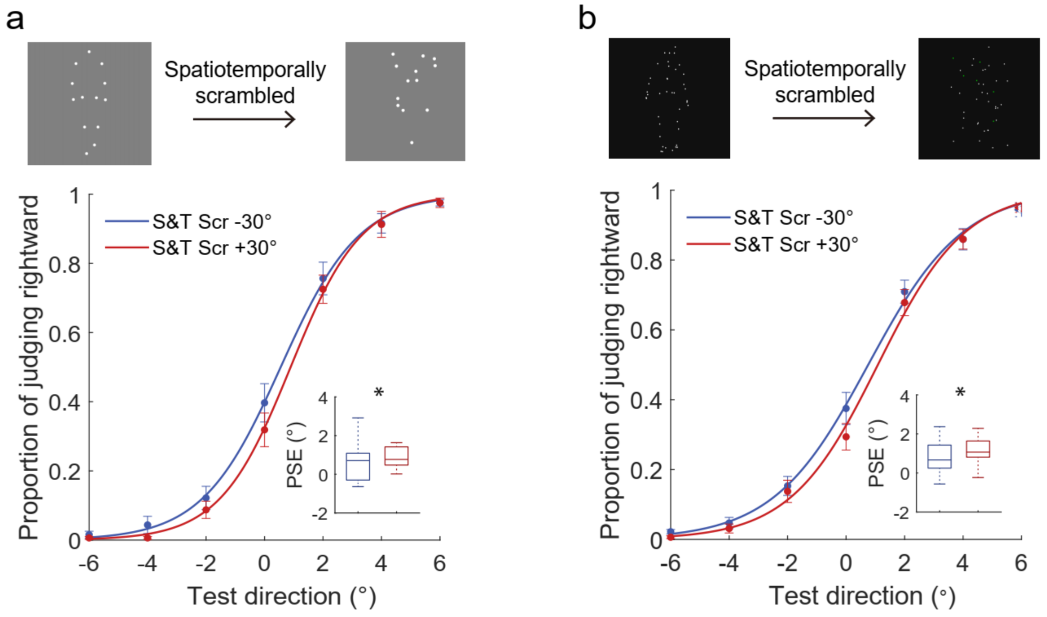


**Figure S1. Results of supplemental experiments.** Stimuli and behavioral data are shown for (**a**) Supplemental Experiment 1 and (**b**) Supplemental Experiment 2. Psychometric functions illustrate viewing direction judgments after adapting to different local BM sequences, and the inset boxplots present the distributions of PSEs for each adapting direction. Asterisks indicate significant differences between the two adapting directions (*$p < 0.05$). "S&T Scr" denotes spatiotemporally scrambled condition.

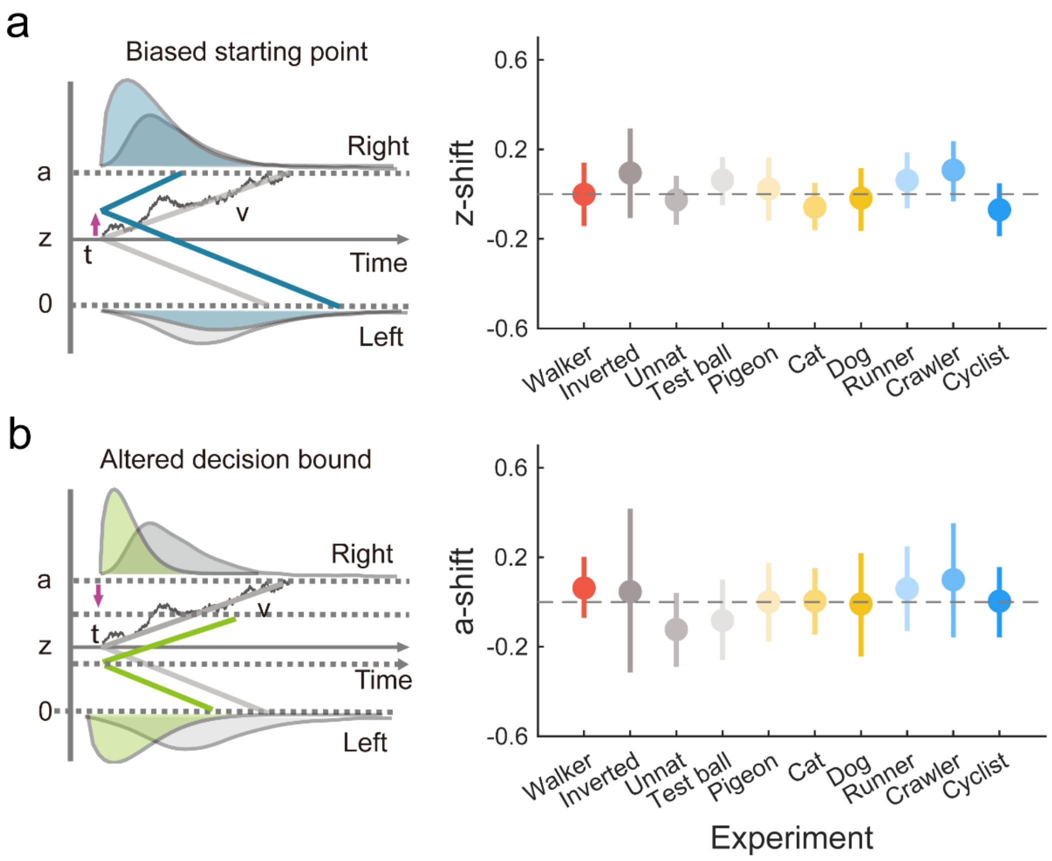


**Figure S2. Drift-diffusion modeling results for decision-level mechanisms.** Schematic illustrations (left) and experimental comparisons (right) for the DDM parameters: (**a**) starting point ($z$, representing prior response bias) and (**b**) decision boundary separation ($a$, representing response caution). The comparisons show adaptation-induced parameter shifts across Experiments 1-3. Error bars indicate the 95% highest density interval of the parameter distribution.

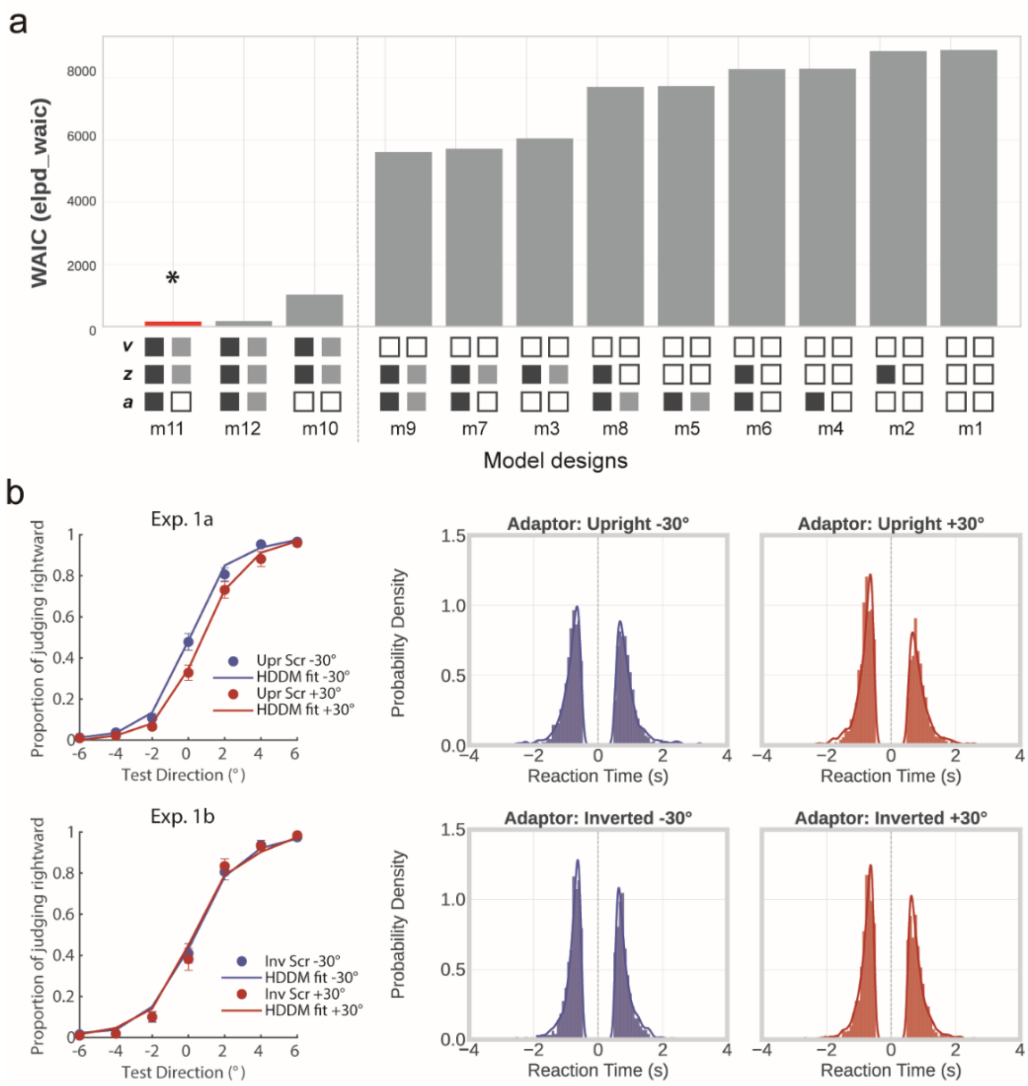


**Figure S3. Model comparison and model simulations using the best-fitting model for Experiment 1a and 1b.** (**a**) Model comparison of 12 candidate drift-diffusion models. Models are ranked by their estimated log predictive density (elpd_WAIC deviance), with lower values indicating better fit. The panel below illustrates model structures: solid squares indicate that the corresponding parameter (*v*, *z*, or *a*) is modulated by the adaptation condition (black) or test stimulus (gray), while empty squares signify that the parameter is not modulated by these factors. Asterisk (*) marks the best-fitting model. (**b**) Model validation through posterior predictive checks. The left panel displays behavioral psychometric functions, and the right panel presents chronometric distributions for left (negative RTs) and right responses (positive RTs). Observed data are shown as blue and red points/histograms for -30° and +30° adaptation conditions. Error bars denote the standard errors of the mean. Model predictions from the best-fitting model (m11), generated from subject-level parameters, are illustrated with blue (-30° adaptation) and red lines (+30° adaptation).

### Supplemental Tables

#### Table S1.

Distribution of adaptation aftereffects and statistical results of paired t tests (two-tailed) comparing PSEs of two adapting directions across experiments.

| | Experimental conditions | PSE shift (°) | | $t$ | $p$ | 95% CI for PSE shifts | | Cohen's $d$ | $BF_{10}$ |
|---|---|---|---|---|---|---|---|---|---|
| | | Mean | SE | | | Lower | Upper | | |
| Exp. 1 | Walker Scr | 0.744 | 0.236 | 3.153 | **0.007**$^{**}$ | 0.241 | 1.247 | **0.788** | **7.746** |
| | Inverted Scr | 0.010 | 0.187 | 0.053 | 0.958 | -0.388 | 0.408 | 0.013 | 0.256 |
| | Unnatural Scr | 0.243 | 0.222 | 1.097 | 0.290 | -0.229 | 0.715 | 0.274 | 0.428 |
| | Test ball | -0.355 | 0.201 | -1.765 | 0.098 | -0.784 | 0.074 | -0.441 | 0.904 |
| Exp. 2 | Pigeon Scr | 0.442 | 0.210 | 2.106 | **0.052**$^{+}$ | -0.005 | 0.890 | **0.526** | **1.448** |
| | Cat Scr | 0.435 | 0.143 | 3.048 | **0.008**$^{**}$ | 0.131 | 0.739 | **0.762** | **6.481** |
| | Dog Scr | 0.764 | 0.209 | 3.657 | **0.002**$^{**}$ | 0.318 | 1.209 | **0.914** | **18.495** |
| Exp. 3 | Runner Scr | 0.451 | 0.215 | 2.092 | **0.054**$^{+}$ | -0.008 | 0.910 | **0.523** | **1.419** |
| | Crawler Scr | 0.452 | 0.113 | 4.007 | **0.001**$^{**}$ | 0.211 | 0.692 | **1.002** | **34.083** |
| | Cyclist Scr | 0.667 | 0.158 | 4.227 | **< 0.001**$^{***}$ | 0.330 | 1.003 | **1.057** | **50.123** |

Note. “Scr” denotes scrambled BMs. Asterisks and numbers in bold indicate significant differences between the two adapting directions (+$p$ = 0.05, **$p$ < 0.01, ***$p$ < 0.001).

**Table S2.**

Summary of statistical results for drift-diffusion modeling parameters across experiments.

| Parameters / Experiments | Drift rate (*v*) | | | Starting point (*z*) | | | Decision bound (*a*) | | |
|---|---|---|---|---|---|---|---|---|---|
| | mean | 95% HDI | $P$ ($L_{30}$ >$R_{30}$) | mean | 95% HDI | $P$ ($L_{30}$>$R_{30}$) | mean | 95% HDI | $P$ ($L_{30}$ > $R_{30}$) |
| Walker Scr | 0.377 | [0.147, 0.591] | **0.998*** | -0.001 | [-0.143, 0.141] | 0.491 | 0.062 | [-0.070, 0.201] | 0.831 |
| Inverted Scr | -0.137 | [-0.458, 0.165] | 0.181 | 0.094 | [-0.107, 0.293] | 0.833 | 0.046 | [-0.314, 0.417] | 0.602 |
| Unnatural Scr | 0.140 | [-0.063, 0.346] | 0.920 | -0.026 | [-0.137, 0.081] | 0.327 | -0.123 | [-0.290, 0.040] | 0.065 |
| Test ball | -0.238 | [-0.447, -0.027] | 0.016 | 0.062 | [-0.048, 0.165] | 0.891 | -0.083 | [-0.258, 0.099] | 0.171 |
| Pigeon Scr | 0.223 | [-0.018, 0.468] | **0.964*** | 0.023 | [-0.119, 0.164] | 0.634 | 0.001 | [-0.176, 0.175] | 0.499 |
| Cat Scr | 0.309 | [0.111, 0.517] | **0.998**** | -0.058 | [-0.161, 0.051] | 0.141 | 0.004 | [-0.145, 0.151] | 0.524 |
| Dog Scr | 0.302 | [0.096, 0.522] | **0.997**** | -0.018 | [-0.164, 0.116] | 0.411 | -0.009 | [-0.243, 0.218] | 0.466 |
| Runner Scr | 0.166 | [-0.054, 0.390] | 0.929 | 0.061 | [-0.064, 0.187] | 0.837 | 0.059 | [-0.129, 0.249] | 0.741 |
| Crawler Scr | 0.141 | [-0.008, 0.297] | **0.967*** | 0.107 | [-0.032, 0.236] | 0.944 | 0.100 | [-0.159, 0.352] | 0.798 |
| Cyclist Scr | 0.264 | [0.061, 0.466] | **0.992**** | -0.070 | [-0.188, 0.048] | 0.123 | 0.004 | [-0.158, 0.156] | 0.528 |

Note. "mean" denotes the mean difference in the parameter distribution between adaptations of +30° and -30°. "95% HDI" refer to the lower and upper bounds of the 95% highest density interval of the parameter distribution. "$P$ ($L_{30}$ >$R_{30}$)" indicates the probability that the given parameter distribution for adapting -30° is greater than that for +30° (*$P$ > 0.95, **$P$ > 0.99).
